# Light and temperature-dependent developmental role of Auxin Binding Protein 1 (ABP1) in Arabidopsis thaliana

**DOI:** 10.1101/2024.01.03.574050

**Authors:** Alena Patnaik, Anshuman Behera, Aman Kumar, Aadishakti Dalai, S Mukundan, Nibedita Priyadarshini, Madhusmita Panigrahy, Kishore CS Panigrahi

## Abstract

Auxin Binding Protein 1 (ABP1) is a small glycoprotein of about 22 kDa that has long been debated as the auxin receptor, and has been put into question for its unclear functions. Despite its conservancy during land plant evolution, its precise role in plant development is still elusive. Historically, it has been implicated in various rapid responses such as membrane polarization, calcium fluxes, TMK1-based cell-surface signalling, auxin canalization, etc. A relatively recent observation questioning the role of ABP1 in plant development led us to explore its probable functions if any. In the current study, we reinvestigated the plausible function of ABP1 using its CRISPR-based loss-of-function mutants, namely *abp1-C1* and *abp1-C2*. Here we show that, ABP1 acts as a positive regulator for primary root elongation under red and secondary root elongation under blue light in seedlings at 22 °C. Under red light at 18 °C, it has a negative effect on hypocotyl growth inhibition. Furthermore, it is involved in flowering time control at 18 °C irrespective of the photoperiod. We show that the transcript levels of Phytochrome B (phyB) and GIGANTEA (GI) are altered in the mutants of ABP1 under red light and low temperature (18 °C) regimes. Further, ABP1 show a pronounced role in tolerance to dehydration induced due to low temperature (18 °C), which correlates with an increase in endogenous abscisic acid (ABA), salicylic acid (SA), and a decrease in jasmonic acid (JA) content in leaves. The functional roles of ABP1 under red light, low temperature and dehydration tolerance in *Arabidopsis thaliana* once again frames it to be an important regulator under adverse and varied conditions that the plant can experience, and thus opened up new avenues for further studies.

## INTRODUCTION

Auxin Binding Protein 1 (ABP1) has been a protein of mystery with its undulating confusions pertaining to its, history (Hertel et al., 1972), and structural properties of having C-terminal KDEL as an ER retention sequence (Tian et al., 1995). ABP1 stood for its name for being crystalized with bound auxin in affinity chromatography assays in many species including maize (Hesse et al., 1986) and tobacco (Shimomura et al., 1999) showing high binding affinities with a synthetic auxin 1-naphthalene acetic acid (NAA) or endogenous auxin (IAA) respectively. Though it has an auxin binding site, neither any conformational change was observed due to bound or unbound auxin nor any response in its transcripts levels due to auxin treatment could be detected. Furthermore, ABP1 was found to be outstandingly stable, not affected by any stimulus, hormone or stress treatment other than heat stress, which leads to its degradation. Notwithstanding, an inconsistent and mild-hiked response of ABP1 to some pathogens was also found (Oliver et al., 1995). Extensive gene expression studies could scarcely find any distinctive profiles of ABP1 except some tissue-specific expressions in the active cambium, embryo tissue, germinating endosperm, stele, hydathodes and trichomes which could hardly explain its dedicated role in auxin perception in plants. ABP1 is speculated with several auxin-induced signalling events requires its expression in the apoplast. However, its ER retention signal sequence which confirms more than 80% of its presence in ER. Even, ABP1-like sequences with or without the KDEL ER retention sequences have also been detected in lower plants (Tromas et al. 2010; Panigrahi et al., 2009). These discrepancies has been explained by the findings that the TIR1 or AFB1, the bonafide auxin receptor is necessarily involved in the rapid polarization events at the plasma membrane and it requires intracellular auxin reinforcing the physiological non-relevance of the ABP1’s auxin binding for these rapid responses. Despite the non-sigmoidal kinetics of auxin binding to ABP, it has a higher and specific binding affinity to the synthetic auxin 1-NAA (more than two-fold) compared to the endogenous auxin (IAA) (Hesse et al., 1989). Further to demonstrate ABP1’s role in plants, several overexpression lines of ABP1 were studied in different instances by Bauly et al. (2018), Čovanová et al. (2013); Grones et al. (2015a) Robert et al. (2010), Chen et al. (2014), which linked its role to activities at plasma membrane such as K^+^ influx to applied auxin, promotion of clathrin-mediated endocytosis of the PIN efflux carriers at the plasma membrane and its definite non-involvement in auxin signalling via the TIR1-Aux/IAA coreceptor complex, thus splitting ABP1 from auxin action. Grones et al. (2015a) showed the essential role of auxin binding pocket of ABP1 in mediating clathrin association with membranes for endocytosis regulation suggesting that auxin binding is central to the function of ABP1 (Grones et al. 2015a). Thus, ABP1 could act as a receptor for rapid auxin responses, such as the activation of plasma membrane H^+/−^ATPases, modulation of cytosolic Ca^2+^ concentrations, TMK1-based cell-surface signalling and auxin canalization (Dahlke et al., 2010, Friml et al., 2022). Concerning the physiological response linked to this interaction, the calcium influx with the auxin-induced calcium-binding interactors of ROPs is associated with root growth (Hazak et al., 2019). The Major bottleneck in understanding its biological function was hindered due to the embryo-lethal effect of existing T-DNA insertional mutants. Relatively recent observation using CRISPR/Cas9 technology that the *abp1* null mutants have no effect on plant development was intriguing. Furthermore, a follow-up work attributed to the second site mutation of the adjacent BSM gene (Michalko et al., 2016, Gao et al., 2015). Moreover, ABP1 was also shown to be directly or indirectly involved in auxin and light signalling, as the application of auxin inhibitor inhibited hypocotyl elongation and gravitropism under R and FR-enriched light in *abp1* mutants but not in wild type (Effendi et al. 2013). ABP1-mediated auxin response was shown to positively affect PhyB signalling for shade (FR-enriched) induced hypocotyl growth inhibition. ABP1 was shown to promote the accumulation of the dephosphorylated form of PhyB required for repressing the gene expression along with the positive regulation of the SCF^TIR1/AFB^ pathway (Grones et al. 2015b). The key role of ABP1 was also shown for shoot growth, leaf venation, cell expansion and division in *Boehmeria nivea* L. (Braun et al. 2008). To explore more on functional and divergent roles of ABP1 in plant development, we conducted scrupulous investigations using two independent CRISPR/Cas9 generated knockout lines of ABP1, namely *abp1-c1* (Gao et al., 2015) and *abp1-c2* (unpublished). Single or combinatorial abiotic stresses were imposed under different monochromatic light and plant phenotypes such as seedling development, flowering time and component of circadian clock genes were studied. We observed effect of *abp1* mutation under low-temperature, dehydration stress and /or under different monochromatic light sources including red, far-red and blue in particular. Here, we discovered a distinct role of ABP1 in establishing a link between photoperiod sensing and circadian clock signalling at lower temperatures and also for dehydration tolerance in *Arabidopsis*.

## MATERIALS AND METHODS

### Plant growth conditions and treatments

Two *abp1* CRISPR-Cas9 generated knockout lines of ABP1, namely *abp1-c1* (Gao et al., 2015) and *abp1-c2* in Col-0 background were used in this study. The *abp1-c1* line harboured a 5-bp deletion in the first exon of ABP1 and resulted in a frameshift with the introduction of a stop codon (Gao et al., 2015). The *abp1-c2* line was received from Professor Klaus Palme’s group, generated using CRISPR technology and was shown to be a functional null for ABP1 (unpublished data). Surface sterilized seeds of *Arabidopsis thaliana* Columbia 0 (Col-0) ecotype, *abp1* mutant lines i.e. *abp1-c1* and *abp1-c2* were sown on square plates with full Murashige and Skoog (MS) medium supplemented with 1 % (w/v) sucrose and 0.8 % (w/v) Phytoagar (pH 5.9) and vernalized at 4 °C for 48 hours (h). Seeds were germinated and grown either at 22 °C or at 18 °C under a long day (LD,16 h light and 8 h dark) or short day (SD,8 h light and 16 h dark) photoperiod conditions and irradiated with different light conditions (dark, white, red, far-red, blue). For exogenous auxin treatment, seeds were directly germinated on MS medium supplemented with 10 mM of 1-Naphthaleneacetic acid (1-NAA, active form of auxin) (Catalog No: PCT0809, Himedia), 10 mM of 2-Naphthaleneacetic acid (2-NAA, inactive form of auxin) (Catalog No: N4002, Sigma) and 20 nM of synthetic auxin 2,4-Dichlorophenoxyacetic acid (2,4-D) (Catalog No: PCT0825, Himedia) and grown for 7 days (d) followed by seedling phenotype analysis. LED chambers (Percival Scientific Inc. Model No. E-30LEDL3) were used with red (30 µmol m^−2^ s^−1^, 660 nm), far-red (30 µmol m^−2^ s^−1^, 730 nm) and blue (30 µmol m^−2^ s^−1^, 460 nm). Simultaneous irradiations of blue and red light were also done using the above two monochromatic light panels. LED chambers were maintained at 22 °C in the LD or SD cycle. Light intensity and wavelength were measured by SpectraPen LM 510 (PSI Instruments, Drásov, Czech Republic). For pot experiments, after 2-d of the vernalisation period, the soil pots containing seeds were transferred to either LD or SD conditions in controlled plant growth chambers (Model No. AR36, Percival, USA) irradiated with white light (50 µmol m^−2^·s^−1^) at 18 °C and 22 °C till the end of the experiment. The flatness of the leaves (leaf epinasty) was determined as discussed earlier by Kozuka et al., 2012. The ratio of the straight-line distance between the two edges of the leaf blade to the actual width of the leaf blade along the curved surface was then determined. For seedling phenotype analysis, 7-d old seedlings were used for scanning as described in (Kumari et al., 2019) using an HP Scanjet G4010 Flatbed scanner. The root and hypocotyl lengths were measured using ImageJ software (https://github.com/imagej/ImageJ). In order to impose dehydration stress, plants were exposed to water withheld (WW) differently at 22 °C or at 18 °C. Plants grown at 22 °C were subjected to WW for 5-d at of 15-d after germination. In parallel, plants grown at 18 °C were subjected to WW for 10-d at 40-d old stage. The difference in the pattern of WW and age of the plant at 22 °C and 18 °C was done due to the slower metabolism rate and longer flowering time in case of low-temperature conditions. Leaves of stressed and unstressed plants were sampled for physiological and biochemical analyses (i.e., relative water content (RWC), leaf temperature analysis, lipid peroxidation assay, carotenoid estimation, enzyme assays and histochemical detection of superoxide radicals).

### Physiological and Biochemical tests

Carotenoid content: Leaf samples (∼25 mg) were incubated with 3 mL of 80% acetone for 48 h at 4 °C in the dark followed by centrifugation at 13000 RPM for 5 mins. The absorbance (A) of supernatants was recorded at 645 nm, 663 nm and 480 nm. Estimation of carotenoids was done according to Manasa et al., (2023).

Relative water content: Leaf segments were excised from the top 4^th^ expanded leaves and fresh weight (FW) was recorded. The segments were floated on deionized water for about 3 h and turgid weight (TW) was recorded. The leaf segments were dried at 60 °C for 24 h and dry weight (DW) was recorded. RWC was calculated using the formula RWC = {(FW-DW) / (TW-DW)} × 100; Where: FW = Fresh weight; TW = turgid weight

Leaf temperature analysis: Plants were used for studying the leaf temperature as described by Patnaik et al., (2023). Briefly, thermal images from plants were captured after keeping them in a dark room with the FLUKE Infra-red Camera (Model No: Ti450 PRO) equidistantly placed 40 cm away. The images were processed using the SMART VIEW software version 4.8.20 and different parts of the plant showing the highest temperature were selected. Temperature from a minimum of 20 plants with 3 biological replicates was analyzed to obtain average leaf temperature data.

Lipid peroxidation assay: Lipid peroxidation was measured according to Heath and Packer (1968). Malondialdehyde (MDA) amount was determined by the thiobarbituric acid (TBA) reaction. Fresh leaf samples (∼100 mg) were homogenized with 2 mL of 0.25% TBA containing 10% trichloroacetic acid (TCA). The homogenate was incubated at 95 ^°^C for 30 minutes (mins) and centrifuged at 10,000g for 10 mins. Absorbance was recorded at 532 nm and nonspecific absorption measured at 600 nm was subtracted.

Histochemical detection of superoxide radical: Nitro-blue tetrazolium (NBT) staining of stressed and unstressed leaves was done to detect the superoxide radical (O2^−^) using the method of Rao and Davis (1999) with some modifications according to Manasa et al (2023). The leaves were immersed in a 10 mM sodium phosphate buffer maintained at pH 7.0 containing 3 mg / mL nitro-blue tetrazolium (NBT) and 10 mM sodium azide (NaN_3_). The immersed leaves were subjected to vacuum infiltration at 80 kPa for 12 h at room temperature in the dark. Acetic acid: glycerol: ethanol (1:1:3 v/v) solution was used to bleach the stained leaves at 100 ⁰C for 30 mins in a water bath. The leaf samples were then stored at 4 °C in a storage solution containing glycerol: ethanol (1:4 v/v). The leaves of the stressed and unstressed plants were imaged using a Carl Zeiss stereo microscope (Model number: Stereo Discovery. V20) equipped with 5.8X magnification and an Axiocam 305 camera to obtain the total area of the NBT stain. The O2^−^ accumulation in the leaves was interpreted in terms of the percentage of stained area. The percentage of stained area in the leaves was calculated using Image J (version: ImageJ 1.53a). Ten different morphologically similar leaves were chosen from five plants of each category to estimate the O2^−^ accumulation. The experiment was repeated thrice with two biological replicates each time.

Antioxidative enzyme assays: For all enzyme assays done for this study (Saha et al 2016), ∼100 mg of leaf extracts were prepared with 2 mL of 100 mM sodium phosphate buffer, pH 7.0 containing 1% PVP (polyvinylpyrrolidone) and 1 mM EDTA (Ethylenediaminetetraacetic acid) on ice. The homogenate was centrifuged at 4 °C for 20 mins at 14000 rpm. The supernatant was used for the enzyme assays.

Catalase (EC 1.11.1.6) activity was measured according to Aebi (1984). Briefly, 0.1 mL of enzyme extract was added to 1 mL of 50 mM sodium phosphate buffer maintained at pH 7.0. The reaction was initiated by adding 0.1 mL of 100 mM hydrogen peroxide (H_2_O_2_). Decrease in absorbance was recorded at intervals of 30 seconds (s) for 2 mins at 240 nm.

Guaiacol peroxidase (POX) activity was measured according to Rao et al. (1996). Briefly, 100 μL of enzyme extract was taken for the assay and 0.7 mL of sodium phosphate buffer, pH-7.0, and 0.1 mL of 1% guaiacol solution were added. 0.1 mL of 10 mM H_2_O_2_ was added to initiate the reaction. The mixture was incubated for 5 mins at room temperature and a change in absorbance was recorded at 470 nm.

Activity of Superoxide dismutase (SOD) activity was determined using 0.1 mL of enzyme extract as described in Panigrahy et al (2019). Briefly, SOD activity was determined by measuring the decrease in the absorbance of blue-colored formazone and O^⋅2−^ at 560 nm.

### RNA extraction and gene expression study

Total RNA was extracted from 7-d-old seedlings grown in different light conditions. Hypocotyl, root samples or 3^rd^ and 4^th^ leaf samples were separately harvested and ∼500 mg of samples were used for total RNA extraction using RNeasy Plant Mini Kit (supplied by Qiagen, Cat #74136). One µg of total RNA was used to obtain cDNA using Reverse Transcription Super-mix (supplied by Bio-Rad, Cat #1708840). For quantitative RT-PCR (qRT-PCR), the Primer Quest tool (Integrated DNA Technologies, Inc., USA) was used to design gene-specific primers used in the qRT-PCR (Supplementary Table S1). All reactions were carried out in hardshell 384-well PCR plates (Bio-Rad, Cat #HSP3805) in Bio-Rad laboratories’ CFX384 Touch^TM^ Real-time detection system, following the manufacturer’s instructions and iTaq^TM^ universal SYBR green super mix as described recently (Patnaik et al., 2023). ACTIN was used to normalize transcript levels. Relative expression levels of the genes were calculated using 2 ^−ΔΔCt^ method (Pfaffl, 2001). The qRT-PCR reactions were done in triplicates from three biological replicates. Data represented as the mean ±SEM.

### Scanning Electron Microscopy

2^nd^ and 4^th^ leaf samples were collected from Col-0 and *abp1* mutant plants at the bolting stage grown on soil under LD conditions at 22 °C. The leaf samples were cleaned properly, placed on glass coverslips with a drop of water and dehydrated at 60 °C for 12 h. The leaves facing upward (adaxial side) were placed on the carbon tape and the surface of the leaf samples was coated with heavy metal platinum prior to imaging (Pathan et al., 2010) using Carl Zeiss Field Emission Scanning Electron Microscopy (Model-ZEISS GeminiSEM 450). An accelerating voltage of 3-5 kV was applied for imaging at different magnifications. The experiment was repeated in three biological replicates.

### Phytohormone estimation

Defence phytohormones (Jasmonic acid, salicylic acid) were estimated at the metabolite estimation facility of NIPGR, New Delhi as described by Patnaik et al., (2023) and Vadassery et al., (2012) with some minor modifications. Briefly, leaf samples (∼300 mg) were harvested at the bolting stage using liquid nitrogen and ground to powder form using tissue lyser (Tissuelyser II, Retsch, Qiagen) at 30 frequency/ s for 2 mins and lyophilised overnight. Nearly 20 mg of lyophilized samples were extracted in 1 mL of methanol. Forty ng mL^−1^ of D6-jasmonic acid (HPC Standards GmbH, Cunners dorf, Germany), 40 ng m L^−1^ of D4-salicylic acid, 40 ng m L^−1^ of D6-abscisic acid, and 8 ng mL^−1^ of jasmonic acid-[13C6] isoleucine conjugate were used as internal standards. The homogenized samples were centrifuged at 14,000 rpm for 20 mins at 4 °C after shaking for 30 mins. The above step is repeated to pool the supernatants, followed by vacuum evaporation and re-suspension in 500 µL of methanol. Samples were analysed on Exion LC (Sciex ®) UHPLC system using formic acid (0.05%) in water as mobile phase A and acetonitrile as mobile phase B. The column and separation protocols were done as described by Patnaik et al (2023).

### Statistical analysis

All the statistical analyses used in this study were performed using GraphPad Prism version 8.0.1. Test of significance was performed to analyse grouped column graphs using two-way ANOVA analysis with multiple comparisons (Tukey and Sidak) and student’s unpaired T-test. Error bars represent the standard error of the mean (SEM).

## RESULTS

In the current study, the probable role of ABP1 in response to different monochromatic lights, low temperature and water withdrawal (WW) was investigated in seedlings and adult plants of *Arabidopsis thaliana*. We conducted the study using two independent CRISPR mutant lines of *abp1* i.e., *abp1-C1* (Gao et a., 2015) and *abp1-C2* (unpublished) in Col-0 background. The null expression of *ABP1* in the above mutant lines was confirmed using q-RT PCR and sequencing analysis (data not shown).

### ABP1 affects seedling root phenotype under red and blue light

Hypocotyl, primary, secondary and lateral root numbers and lengths were measured as described in materials and method (Section-2.1) at 22 °C under diurnal LD conditions with Red (R, 30 µmol m^−2^ s^−1^ 660 nm)/ far-red (FR, 30 µmol m^−2^ s^−1^, 730 nm)/ blue (B, 30 µmol m^−2^ s^−1^, 460 nm)/ white light (W, 50 µmol m^−2^·s^−1^) or in Dark (D, 0 µmol m^−2^·s^−1^). Among all the light qualities, hypocotyl length and root patterning in *abp1* mutants (i.e., *abp1-C1* and *abp1-C2*) showed a distinct difference as compared to Col-0 when grown at 22 °C under R and B. The *abp1-C1* and *abp1-C2* had considerably shorter hypocotyls (2.8-fold) and roots (1.2-fold) as compared to Col-0 specifically under R light (Figure 1A & 1B), with no significant differences under the rest of the monochromatic lights including FR, B and W light or D (Figure S1). Hypocotyl and root lengths under W light were exactly similar to that shown by Gao et al (2015). Further, the secondary root phenotype (i.e., number of lateral and adventitious roots) in *abp1* mutants under monochromatic R (Figure 1C) or B (Figure 1D) or simultaneous irradiations of R and B (R+B) (Figure 1E) lights were noted. Under R, the secondary root counts were insignificant in all three lines (Figure 1E). In contrast, under diurnal B, the secondary root counts in *abp1* mutants were significantly decreased in the ABP1 mutants (∼70% in *abp1-C1* and ∼65% in *abp1-C2*) as compared to the Col-0 ecotype (Figure 1D). lateral root number. However, the secondary root counts under diurnal R+B light showed marginally significant differences when compared between WT and mutants (Figure 1E), leading to the inference that the decreased secondary root number under B could be effectively overridden by the simultaneous irradiation of B with the R light (i.e., B+R lights). To understand more about R light signalling and the circadian clock, the relative transcript levels of *PHYTOCHROME B* (*PHYB*) and *GIGENTEA* (*GI*) were examined under R light irradiations. There were ∼26-fold and 20-fold increase in the *PhyB* transcript levels in *abp1-C1* and *abp1-C2* mutants respectively as compared to Col-0 (Figure 1F). The relative transcript levels of *GI* in *abp1-C1* and *abp1-C2* were ∼3.8-fold and 4-fold lower than Col-0 respectively (Figure 1G). These results indicated that ABP1 most likely has a role in root development under R light and secondary root growth under B light, with the former role could be mediated by phytochrome B and/or GI.

**Figure 1.**
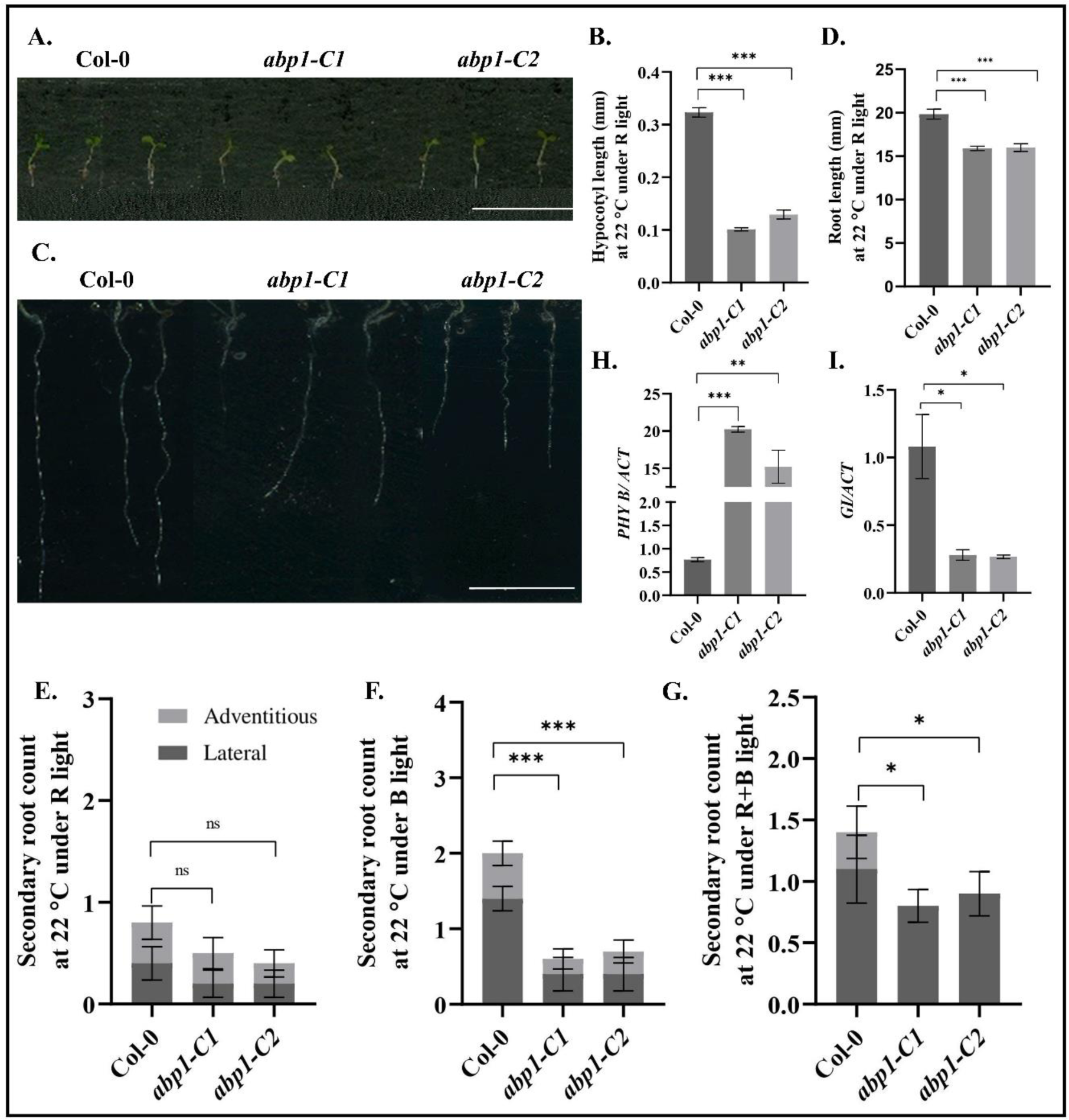
Seedling phenotype under diurnal monochromatic red light at 22 °C. Seedlings of Col-0, *abp1-C1* and *abp1-C2* lines were grown on MS media under monochromatic red light at 22 °C in long-day conditions for 7 d **(A-C),** after which the phenotypes hypocotyl lengths (**B**) root length **(D)** and the number of secondary roots (lateral and adventitious roots) were noted (**E**). Secondary root count was noted for seedlings grown under diurnal monochromatic blue (**F),** or under a simultaneous irradiation of red and blue light **(G)**. Roots from the seedlings grown under red were analysed for transcript expression of Phytochrome B *(PHYB)* (**H)** and *GIGANTEA (GI) (***I)** using qRT-PCR. For hypocotyl and root phenotype, 300 seedlings were grown in 3 biological replicates with 100 seedlings each time. The qRT-PCR reactions were done in triplicates from three biological replicates. *ACTIN* was used to normalize transcript levels. Data presented as mean ± SE. One-way analysis of variance (ANOVA) using Tukey’s multiple comparisons test was performed with the help of GraphPad Prism to test for significance among the dataset, * *p* < 0.05, ** < 0.01, *** *p* < 0.001and ns represent non-significant differences. Scale bar: 1 cm.

### ABP1 mediates hypocotyl elongation under diurnal red rhythm along with low temperature

In addition to the hypocotyl and root phenotype at the seeding stage at 22 °C, it was of interest to check the seedling phenotype if any with effect to low temperature (i.e., 18 °C). Congruent with the data of Gao et al 2015, no remarkable difference in the hypocotyl and root length was observed under 22 °C under W light (Figure S1). Similarly, no phenotypic difference was observed at 18 °C in *abp1* mutants compared to Col-0 under W light irrespective of photoperiod i.e., LD (Figure S2A-S2C) or SD (Figure S2D-S2F) conditions. However, the hypocotyl lengths of *abp1* mutants were significantly different at 18 °C under diurnal R light both under LD (Figure 2A) and SD (Figure S2D-S2F) photoperiod. The hypocotyl length of 7-d-old *abp1* mutant seedlings grown in LD conditions was significantly shorter (1.36-fold in *abp1-C1* and 2.08-fold in *abp1-C2*) than Col-0 wild type (Figure 2E & 2F). Similarly, the *abp1* mutants (1.2-fold in *abp1-C1* and 1.2-fold in *abp1-C2*) showed shorter hypocotyl with respect to Col-0 lines under SD conditions at 18 °C (Figure S2G-S2I). In addition, the root lengths of *abp1* mutants were marginally shorter than the Col-0 under R light at 18 °C irrespective of the LD or SD photoperiods (Figure S2K & S2L). The transcript levels of *PHY B* were 1.39-fold and 1.58-fold higher in *abp1-C1* and *abp1-C2* mutants respectively than that of Col-0 at 18 °C (Figure 2G). The transcript level of *GI* was significantly decreased (i.e., 2.48-fold) in the *abp1-C1* than in Col-0 wild-type plants. Though the decrease in *GI* transcript level in *abp1-C2* was not as severe as in *abp1-C1* still it was significant (p ≤ 0.05) (Figure 2H). These results indicated that ABP1 is involved in hypocotyl growth inhibition under monochromatic R light at lower temperature. Downregulation in *GI* gene expression also suggested that these seedling phenotypic changes in *abp1* mutants are directly or indirectly GI-mediated. Moreover, the role of ABP1 in controlling the seedling phenotype (hypocotyl length and root growth) under low temperature appears to have a conserved yet unidentified mechanistic function to that of ambient temperature (i.e., 22 °C).

**Figure 2.**
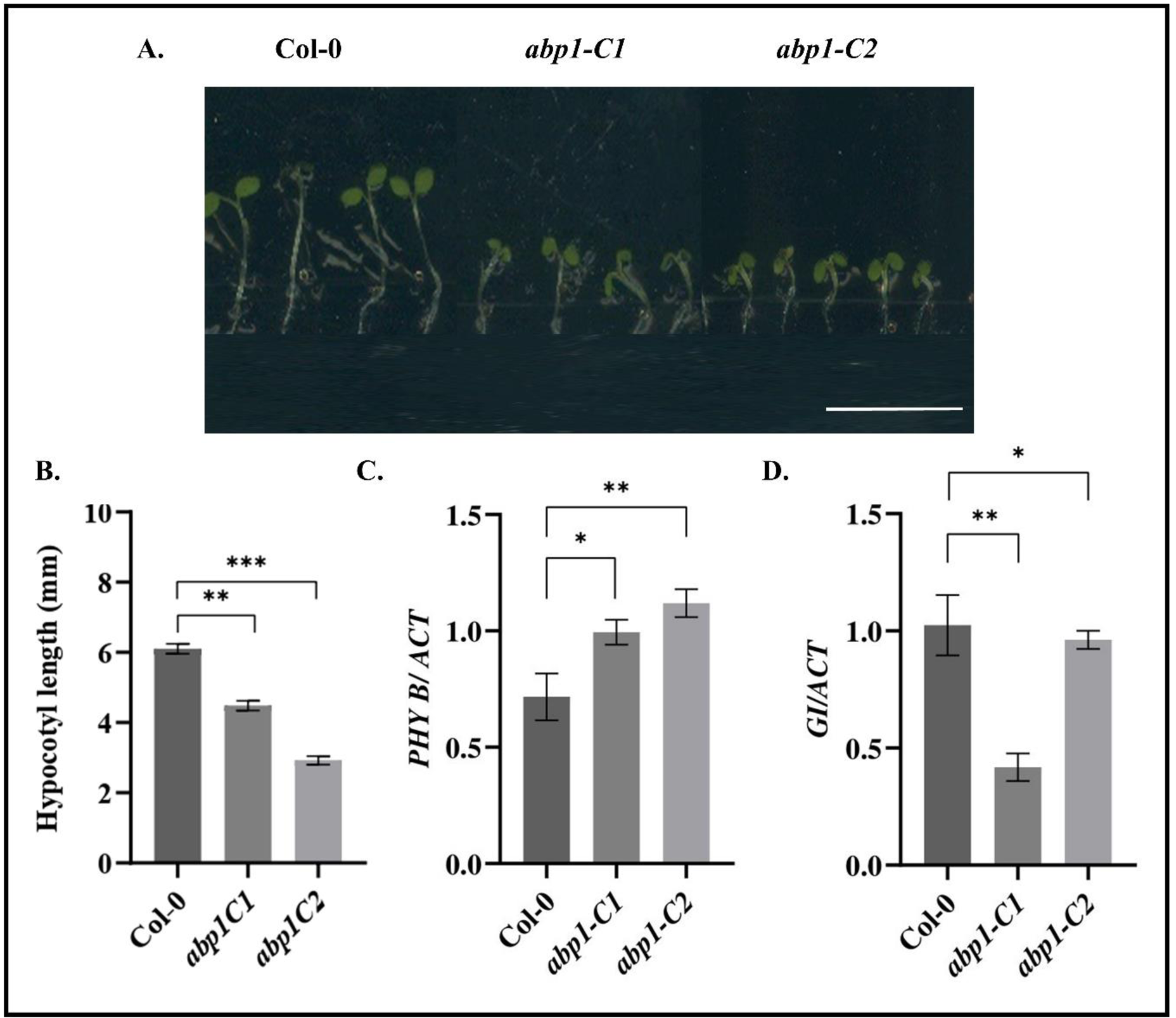
Hypocotyl growth inhibition under monochromatic red light and lower temperature (18 °C). **A.** Seedlings of Col-0, *abp1-C1* and *abp1-C2* genotypes were grown on MS media under monochromatic red light at 22 °C and long-day conditions for 7-days (**A**), after which hypocotyl length was measured (**B).** Hypocotyl from the seedlings grown under red was analysed for transcript expression of Phytochrome B *(PHYB)* (**C)** and *GIGANTEA (GI)* (**D).** The hypocotyl lengths were measured using ImageJ software. Data presented as mean ± SE. The qRT-PCR reactions were done in triplicate from three biological replicates. *ACTIN* was used to normalize transcript levels. One-way analysis of variance (ANOVA) using Tukey’s multiple comparisons test was performed with the help of GraphPad Prism to test for significance among the dataset, * *p* < 0.05, ** < 0.01, *** *p* < 0.001and ns represent non-significant differences. Scale bar: 1 cm.

**Figure 3.**
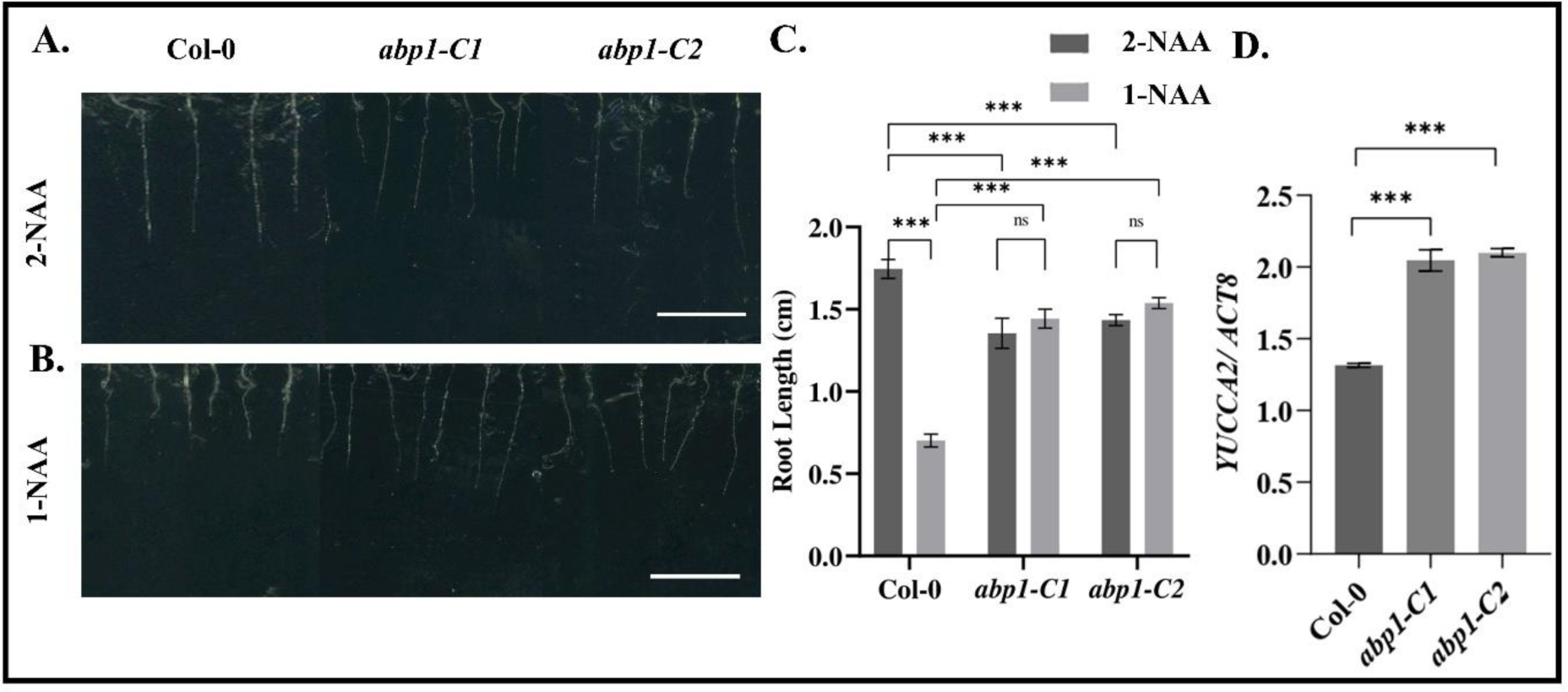
Complementation of root growth phenotype with external application of synthetic auxin. Seedlings of Col-0, *abp1-C1* and *abp1-C2* genotypes were grown on MS media under monochromatic red light, 22 °C and long-day conditions for 7-days and f root phenotype was studied after supplement of 1 mM 1-naphthaleneacetic acid (NAA) active form of auxin (**B**) and 2-NAA, inactive form of auxin **(A)** or with 1 mM 1-NAA, after which root length were measured **(C).** The 3^rd^ and 4^th^ leaves of adult plants (10-d before bolting) were used for analyses of transcript expression of *YUCCA2* (**D**). 300 seedlings were grown in 3 biological replicates with 100 seedlings each time. The root lengths were measured using ImageJ software. Data presented as mean ± SE. One-way analysis of variance (ANOVA) using Tukey’s multiple comparisons test was performed with the help of GraphPad Prism to test for significance among the dataset, * *p* < 0.05, ** < 0.01, *** *p* < 0.001and ns represent non-significant differences. Scale bar: 1 cm. D-Expression of *YUCAA2* (adult leaves) fold change-The levels of YUCCA2 genes in *abp1-C1* and *abp1-C2* increased by 1.5 and 1.5-fold respectively as compared to Col-0.

### Synthetic auxin 1-NAA can rescue abp1 mutant root phenotype under red light

Since *abp1-C1* and *abp1-C2* mutants showed R light-specific root phenotype strikingly under 22 °C (Figure 1) and marginally under 18 °C (Figure S2K & S2L), a rescue assay was performed using synthetic auxins i.e.,1-NAA (active form of auxin) and 2-NAA (inactive form of auxin). Col-0 lines showed a 51-57 % decrease in root length with 10 mM 1-NAA supplement in the growth medium (Figure 2). In both the *abp1-C1* and *abp1-C2* lines, the root length attained to those of their respective controls with 10 mM of 2-NAA in the media (Figure 2). A similar but less dramatic observation was also obtained when another synthetic auxin i.e., 2,4-dichlorophenoxyacetic acid (2,4-D) was used (data not shown). These results indicated that while exogenous auxin supplement resulted in root length inhibition in Col-0, it could complement the growth inhibition in the absence of ABP1 (in the *abp1* mutant lines). Hence, it led to the explanation that the decrease in the root length in Col-0 under R light was very likely due to ABP1 binding to auxin in the roots. These results convincingly suggested that ABP1 has an important role in root development under R light (660 nm) of 30 µmol m^−2^ s^−1^ fluence and pronounced at 22 °C growth conditions while compared to that of 18 °C.

### ABP1 mutation delays timing to flower at low temperature irrespective of photoperiod

Since *abp1* mutants showed higher *PHYB* and lower *GI* transcript levels at lower temperature (Figure 2), the adult plant phenotypes in the above conditions were intrigued. Both the CRISPR mutants of ABP1 under W light at 22 °C showed indistinguishable phenotypes from that of the Col-0, which was similar to results obtained in the case of *abp1-C1* by Gao et al. (2015) (Figure S3). However, both the *abp1-C1* and *abp1-C2* mutants showed delayed flowering with an increased number of rosette leaves under W light at 18 °C irrespective of photoperiod (i.e., under both 16/8h, LD and 8/16h, SD) (Figure 4A and Figure 4C). Explicitly, the *abp1-C1* and *abp1-C2* mutants had a significantly higher number of rosette leaves, 23±2 and 25±2 respectively than the Col-0 with 16±2 rosette leaves at the time of bolting under LD conditions (Figure 4B). Similarly, under SD conditions the *abp1-C1* and *abp1-C2* mutants had 59±2 and 57.8±2 leaves respectively compared to 45±2 rosette leaves in Col-0 at the time of bolting (Figure 4D). Since the delayed flowering phenotype of *abp1* mutants under low temperature was irrespective of LD or SD photoperiodic conditions, all further experiments were carried out under LD conditions. Rosette radius in *abp1-C1* and *abp1-C2* mutants also increased to 1.1-fold and 1.2-fold respectively as compared to the Col-0 lines (Figure 4E). Transcript expression levels of *FT* in Col-0 in comparison to those of *abp1* mutants grown at 18 °C were studied to explain the above-mentioned flowering phenotype. The expression levels of *FT* in *abp1-C1* and *abp1-C2* were observed to be 12.3-fold and 1.69-fold lower respectively than Col-0 (Figure 4F). Previous reports showed a larger rosette radius in the late flowering mutants like *gi* (Weraduwage *et al.,* 2015). Since the rosette radius was larger in the mutants, the expression of *GI* in *abp1* mutants was investigated. The *GI* transcript level was found to be significantly lower (i.e., 6.48-fold and 3.91-fold in *abp1-C1* and *abp1-C2* respectively) than Col-0 (Figure 4G).

**Figure 4.**
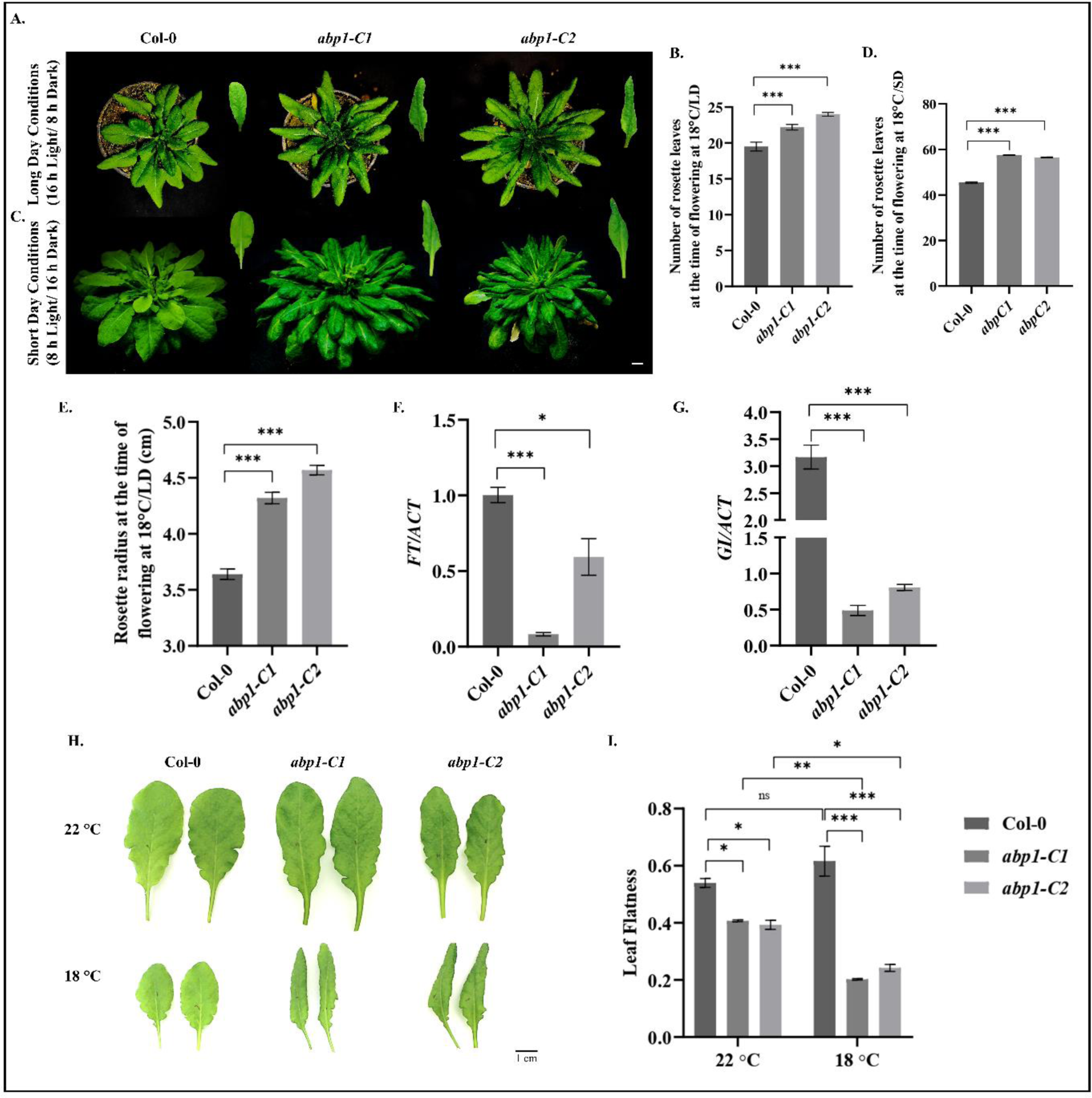
Flowering phenotype at lower temperature (18 °C). Plants of Col-0, *abp1-C1* and *abp1-C2* genotypes were grown on soil in pots under LD (**A-B**) and SD (**C-D**) conditions and 18 °C till bolting. Rosette leaf number (**B & D)** and radius at the time of flowering (**E**) were noted. Three biological replicates were performed using 10 plants in each replicate. The 3^rd^ and 4^th^ leaves were used for analyses of transcript expression of *FLOWERING TIME* (*FT)* (**D)** and *GIGANTEA (GI)* (**E**). Representative leaves from each genotype are shown to show epinasty (**H**) and histogram representing the leaf flatness in each genotype at 22 °C and 18 °C. Data presented as mean ± SE. The qRT-PCR reactions were done in triplicates from three biological replicates. *ACTIN* was used to normalize transcript levels. One-way analysis of variance (ANOVA) using Tukey’s multiple comparisons test was performed with the help of GraphPad Prism to test for significance among the dataset, * *p* < 0.05, ** < 0.01, *** *p* < 0.001and ns represent non-significant differences. Scale bar: 1 cm.

Another remarkable phenotype in *abp1* mutants was leaf epinasty. The leaves of ABP1 mutants were found to be more curled towards the abaxial surfaces (Figure 4H). The leaf flatness is expressed as the ratio of the straight-line distance between the two leaf edges to the actual leaf width (Kozuka et al., 2012). The leaves of *abp1-C1* and *abp1-C2* were 3.04-fold and 2.56-fold more curled than the Col-0 leaves at 18 °C (Figure 4I). However, the leaves of *abp1* mutants were found to be completely expanded with a marginal difference at 22 °C (Figure 4I). The above results convincingly showed that *abp1* mutation delays flowering and causes leaf epinasty at low temperature.

### Role of defence hormones in mediating low-temperature response in abp1 mutant lines

Cold responses and acclimatization are known to be associated with the dynamic interplay of several hormones including jasmonic acid (JA) (Wang et al., 2020), abscisic acid (ABA) and salicylic acid (SA) in plants (Shi and Yang, 2014; Scott et al 2004). As *abp1* mutants showed prolonged growth effects with a higher number of leaves and delayed flowering under lower temperature, we quantified the levels of endogenous hormones such as ABA, SA and JA under lower temperature in *abp1* mutants. While Col-0 showed a 13.55 % decrease under 18 °C, *abp1-C1* and *abp1-C2* lines had 28.35 % and 6.56 % increase in ABA content respectively when compared with their respective controls under 22 °C (Figure 5A). Similarly, Col-0 lines showed a 9.08 % decrease, whereas *abp1-C1* and *abp1-C2* lines had 20.46 % and 22.90 % increase in SA content respectively in plants grown under 18 °C (Figure 5B). On the contrary, Jasmonic acid (JA) content showed a reverse trend of a 2.37-fold increase in response to lower temperature in Col-0 lines. The *abp1-C1* and *abp1-C2* lines exhibited a 61.92 % and 76.60 % decrease in JA content in similar conditions (Figure 5C). These results indicated a correlation between the sub-lethal growth phenotype under reduced temperature in the *abp1* mutants with the accumulation of dormancy and defence hormones such as ABA and SA.

**Figure 5.**
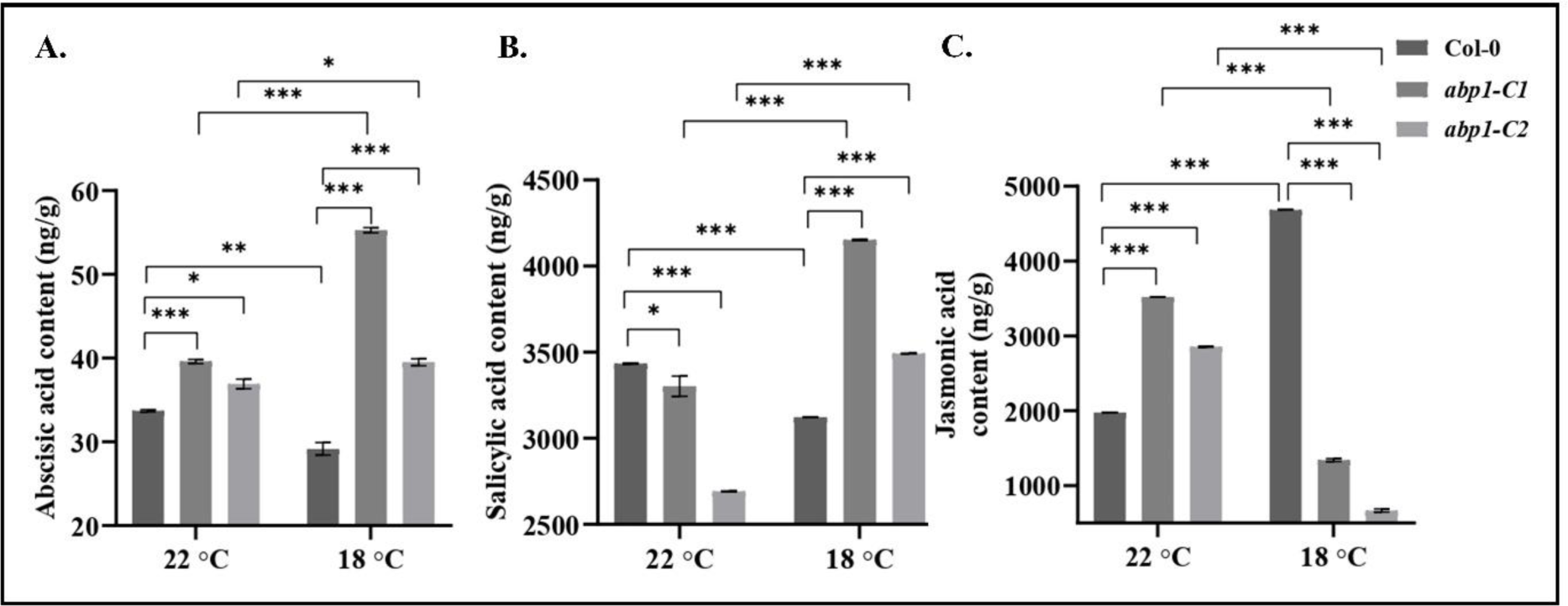
Estimation of endogenous contents of defense hormones (abscisic acid, salicylic acid and jasmonic acid). Leaf samples from adult plants grown on soil in pots under LD conditions from Col-0, *abp1-C1* and *abp1-C2* lines grown under either 22 °C or 18 °C were collected at bolting stage, which were processed for determination of endogenous contents of abscisic acid (ABA) (**A),** Salicylic acid (SA) (**B**) and jasmonic acid (JA) (**C**). D6-ABA, D4-SA and D6-JA were used as internal standards for ABA, SA and JA quantification. Three individual biological replicates were performed to obtain the mean ± SE. One-way analysis of variance (ANOVA) using Tukey’s multiple comparisons test was performed with the help of GraphPad Prism to test for significance among the dataset, * *p* < 0.05, ** < 0.01, *** *p* < 0.001.

### Altered trichome morphology contributes to cold resistance in abp1 mutants

As trichomes are known to be involved in several abiotic stresses (Wang et al., 2021) including cold tolerance, trichome density and architecture were studied in plant leaves grown under W light. As shown in previous studies, low-temperature stress resulted in varied alterations (either lower or higher) in trichome density (Sun et al. 2022; Zhang et al., 2020) across different plant species. In the Col-0 line trichome density was increased by 2.9-fold under lower temperature (Figure 6A & 6B). However, in *abp1-C1* and *abp1-C2* lines the trichome density changed insignificantly at 18 °C compared to their respective controls at 22 °C leading to reduced trichome density than those of Col-0 at 18 °C (Figure 6A & 6B). Trichome architecture and branching have also been known to vary in response to abiotic stresses (Iordachescu et al., 2008). A number of genes regulate the secondary branching of trichomes, either positively or negatively (Zhang et al., 2021). Secondary trichome branching was not observed in Col-0 lines both under control and low temperatures conditions. However, secondary branching (i.e., four branched trichomes) increased by 58.3 % and 38.7% in *abp1-C1* and *abp1-C2* lines respectively at 22 °C as compared to those of Col-0 (Figure 6C & 6D). In contrast, under low temperature, 2.75-fold and a marginal decrease in secondary branching were observed in *abp1-C1* and *abp1-C2* lines (Figure 6C & 6D) compared to their controls at 22 °C respectively. TRIPTYCHON (TRY) acts as a negative regulator of secondary branching of trichomes and its loss of function results in the formation of new branch points (Lescot et al., 2002). CAPRICE (CPC) is a negative regulator of trichome initiation and development (Fambrini et al., 2019). The trichome branch pattern in the *abp1* mutants was validated genetically by analysing the transcript levels of *CPC* and *TRY* genes. Under control growth conditions transcript levels of *CPC* were 12.9-fold and 11-fold less in the *abp1-C1* and *abp1-C2* lines respectively than in the Col-0 (Figure 6E). Similarly, the transcript levels of *TRY* were 3-fold and 1.4-fold less in *abp1-C1* and *abp1-C2* lines respectively than the Col-0 (Figure 6F). With the effect of low-temperature stress, *CPC* and *TRY* transcript levels increased non-significantly, in both *abp1-C1* and *abp1-C2* lines, when compared to their respective controls grown at 22 °C (Figure 6E & 6F). These results suggested that phenotype in *abp1* mutant lines is associated with decreasing trichome density and increasing branching patterns which in turn may directly or indirectly be regulated by an increase in transcript levels of negative regulators of trichome branching genes such as *CPC* and *TRY*.

**Figure 6.**
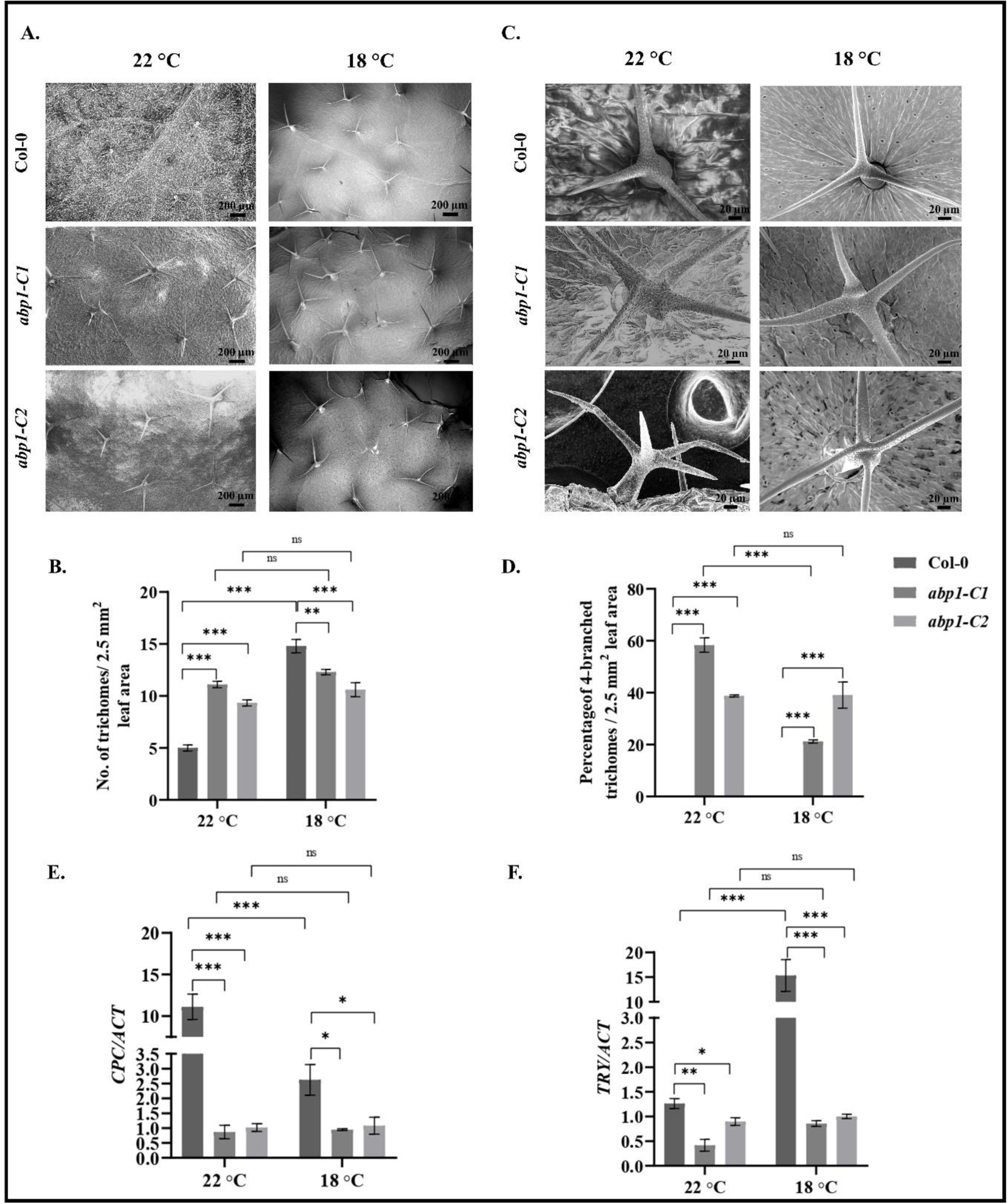
Molecular analysis of trichome morphology. Plants of Col-0, *abp1-C1* and *abp1-C2* genotypes were grown on soil in pots under LD conditions at 22 °C and 18 °C till bolting followed by leaf sample collection. Leaf samples were studied using scanning electron microscopy (SEM) (**A & C**) as described in methods section. Trichome density (**B**) and percentage of 4-branched trichomes (**D**) were analysed from SEM micrographs. Percentage of 4-branched trichomes were calculated to the total number of trichomes (**D**). Three individual biological replicates were performed. Data presented as mean ± SE. Leaf samples were used for transcript expression analysis of *CPC* **(E)** or *TRY* (**F**). Reactions were done in triplicate from three biological replicates. *ACTIN* was used to normalize transcript levels. One-way analysis of variance (ANOVA) using Tukey’s multiple comparisons test was performed with the help of GraphPad Prism to test for significance among the dataset, * *p* < 0.05, ** < 0.01, *** *p* < 0.001.

### ABP1 mutation positively affects dehydration tolerance

Cold or drought stress induces dehydration is well documented (Bhat et al. 2022, Manasa et al. 2022, Zhang et al., 2018). Divulged of the cold resistance effect due to ABP1 mutation, we investigated the effect of dehydration induced by water withheld (WW) at 22 °C or at low temperature (18 °C) in *abp1* mutant lines (Figure 7A).

**Figure 7.**
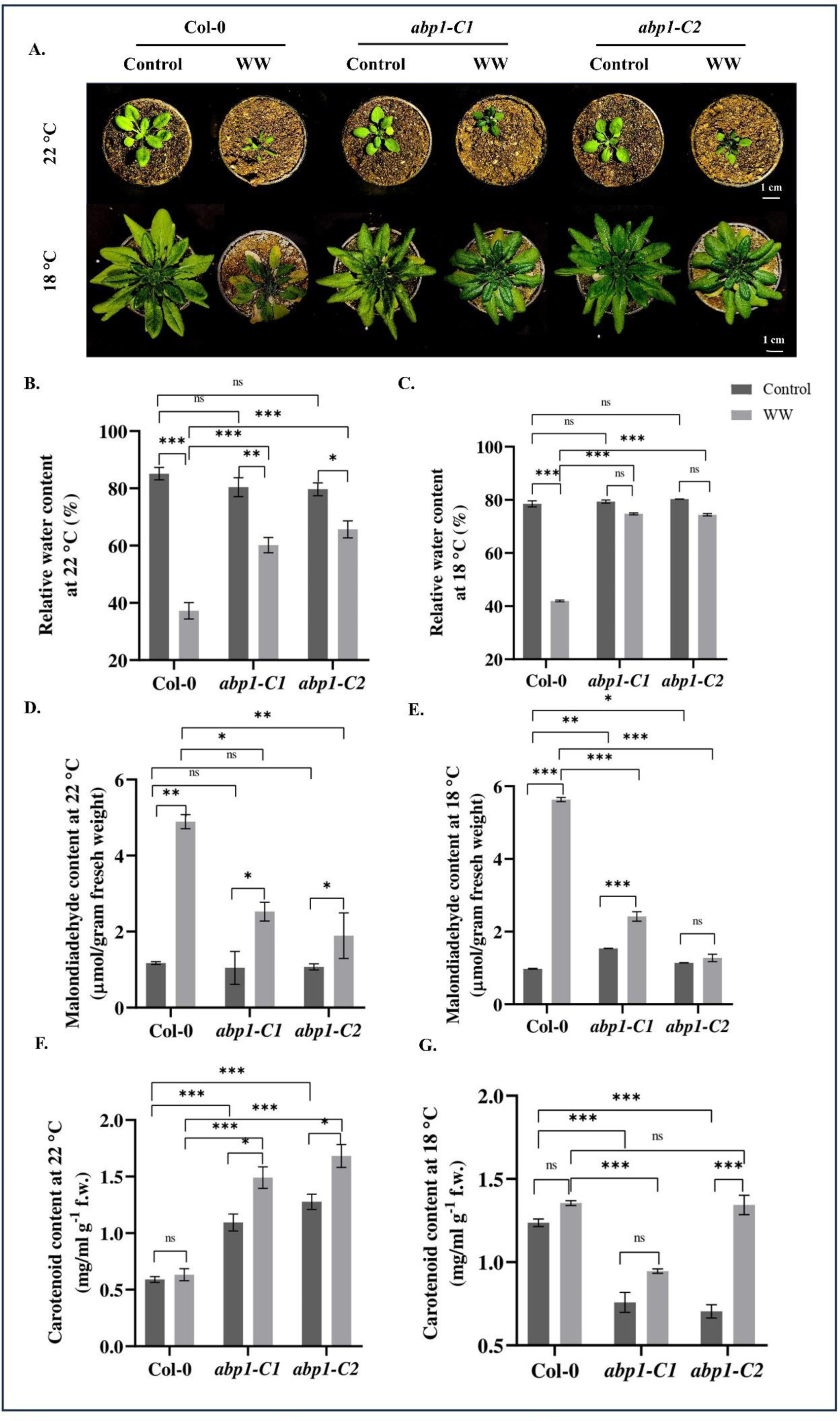
Analysis dehydration stress tolerance. Plants of Col-0, *abp1-C1* and *abp1-C2* genotypes were grown on soil in pots under LD conditions at 22 °C or 18 °C till 15-d or 40-d old stage respectively (as described in section 2.1). Water was withheld (WW) for 5-d at 22 °C (**A, top row**) or 10-d at lower temperature (**A, bottom row**) were imposed to induce dehydration stress. Physiological parameters including relative water content (**B**, **C**), malondialdehyde content (**D**, **E**) and total carotenoid content (**F**, **G**) were studied from the leaf samples subjected to WW at 22 °C and 18 °C respectively. Three individual biological replicates were performed. Data presented as mean ± SE. One-way analysis of variance (ANOVA) using Tukey’s multiple comparisons test was performed with the help of GraphPad Prism to test for significance among the dataset, \**p* < 0.05, **< 0.01, \*\*\**p* < 0.001, ns: non-significant. Scale bar: 1 cm.

#### Physiological analysis of dehydration tolerance

Relative water content (RWC) decreased to nearly half of its initial value due to WW in Col-0 under both 22 °C (Figure 7B) and low temperature (Figure 7C), whereas in *abp1-C1* there was only a minimal decrease in the RWC percentage (∼25.1 % at 22 °C and marginal decrease of about 5.7 % at 18 °C) after 5d or 10d of WW respectively. The *abp1-C2* also showed an identical pattern of decrease in RWC as that of *abp1-C1* (Figure 7B & 7C). Malondialdehyde (MDA) accumulation due to lipid peroxidation in plant leaves serves as an exclusive indicator of abiotic stress response (Singh et al., 2009; Manasa et al., 2023). Col-0 plants showed a significant increase of MDA content after WW i.e., 4.1-fold and 5.8-fold as compared to their respective controls at 22 °C as well as under 18 °C respectively (Figure 7D & 7E). However, the MDA accumulation after WW in *abp1-C1* and *C2* lines was significantly less as compared to that of Col-0. Moreover, both *abp1-C1* and *abp1-C2* showed similar patterns of MDA accumulation, i.e., nearly 2.4-fold and 1.76-fold (at 22 °C) (Figure 7D) as well as 1.5-fold and 1.1-fold (at 18 °C) (Figure 7E) increase than their respective control plants. The endogenous temperature of plants after WW concerning their respective controls increased slightly at both 22 and 18 °C (Figure S4).

#### Biochemical analysis of dehydration tolerance

Dehydration due to drought or cold is associated with characteristic physiological changes including accumulation of stress pigments such as carotenoids and flavonoids, reduction of photosynthetic efficiency, generation of reactive oxygen species (ROS) and increased ROS scavenging due to various ROS scavenging enzyme activity intolerant species, which have been well documented in previous studies (Panigrahy et al 2011, Tomanek et al 2012, Manasa et al 2022, Panigrahy et al 2022). Dehydration tolerance in *abp1* mutant lines was validated after WW by analysing various biochemical parameters such as carotenoid content, oxygen free radicals (O2^.-^) accumulation, superoxide dismutase (SOD), catalase (CAT) and peroxidase (POX) activities in comparison with those of Col-0 as well as with their respective controls without WW.

Carotenoid content due to WW at both 22 °C and 18 °C was found to be negligible in the Col-0 line while both the *abp1* lines showed higher carotenoid accumulation after WW (i.e., 1.3-fold, 1.3-fold higher carotenoids at 22 °C (Figure 7F) and 1.25-fold, 1.9-fold higher carotenoids at 18 °C (Figure 7 G) in *abp1-C1* and *abp1-C2* line respectively when compared to their corresponding controls)

Col-0 lines showed a sharp visible increase in accumulation of O2^.-^ due to WW as evident from the histochemical staining of the leaves (3.2-fold and 2.5-fold increase) both at 22 °C (Figure 8A & 8B) and 18 °C (Figure 8C & 8D) respectively. Dehydration stress due to WW showed significantly less O2^.-^ accumulation in both the *abp1* lines as compared to that of Col-0 at 22 °C (Figure 8B). Explicitly, while *abp1-C1* showed a 1.7-fold increase, the *abp1-C2* line showed a 1.7-fold increase in O2^.-^ respectively at 22 °C (Figure 8B). Under 18 °C, both the *abp1* lines showed a negligible or insignificant increase in O2^.-^ as compared to their respective controls (Figure 8D).

**Figure 8.**
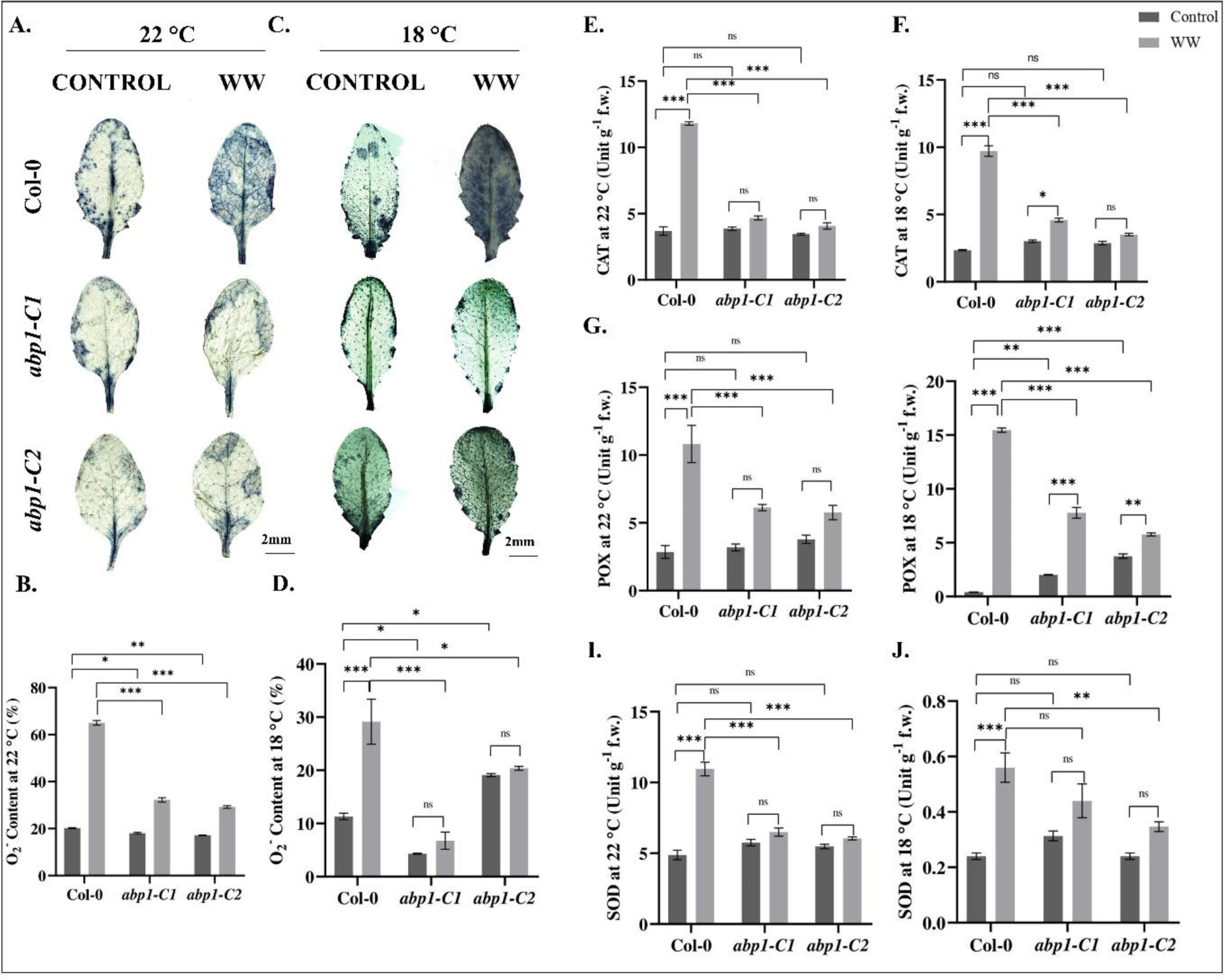
Biochemical analyses during dehydration stress. Plants of Col-0, *abp1-C1* and *abp1-C2* genotypes were grown on soil in pots under LD conditions at 22 °C or 18 °C till 15-d or 40-d old stage respectively (as described in section 2.1). Water was withheld (WW) for 5-d at 22 °C or 10-d at lower temperature were imposed to induce dehydration stress. Histochemical detection of oxygen radical at control, 22 °C (**A, B**) and 18 °C (**C, D**) was done using NBT staining (**A, C**) and percentage of oxygen radical (O^⋅2−^) accumulation (**B, D**) in the leaves by calculating the stained area to the total area of the leaves using ImageJ software. Determination of ROS scavenging enzyme activity including catalase (CAT) (**E, F)**, Guaiacol peroxidase (POX) (**G, H**) and Superoxide dismutase (SOD) activity (**I, J**) under control and low temperature were done using spectrophotometric assays. Three individual biological replicates were performed for each experiment with 3 plants each sample. Data presented as mean ± SE. One-way analysis of variance (ANOVA) using Tukey’s multiple comparisons test was performed with the help of GraphPad Prism to test for significance among the dataset, \**p* < 0.05, **< 0.01, \*\*\**p* < 0.001, ns: non-significant. Scale bar: 2 mm.

WW at both 22 °C and 18 °C resulted in a significant increase in catalase (CAT), peroxidase (POX), and superoxide dismutase (SOD) enzyme activities in Col-0 plants. At 22 °C, the increases were notable, with a 3.2-fold, 3.8-fold, and 2.2-fold rise in CAT, POX, and SOD activities, respectively (Figure 8E, 8G & 8I). Similarly, at 18 °C, the enzyme activities in Col-0 plants exhibited substantial increments to 4.14-fold, 38.6-fold and about 2.3-fold for CAT, POX, and SOD, respectively (Figure 8F, 8H & 8J). However, in comparison to Col-0, the *abp1* mutant lines showed relatively smaller insignificant increase in enzyme activities after WW. However, close observation of the *abp1* lines in detail, after WW at 22 °C, there was a 1.9-fold and 1.5-fold increase in POX activity in *abp1-C1* and *abp1-C2* lines, respectively. Similarly, at 18 °C, an increase of 1.5-fold, 1.2-fold in CAT activity, 3.8-fold, 1.5-fold in POX activity and 1.4-fold, 1.4-fold in SOD activity was observed in *abp1-C1* and *abp1-C2* lines, respectively, when compared to their respective control conditions.

## DISCUSSION

Our results demonstrated for the first time a conditional role of ABP1 predominantly under R light and lower temperature (18 °C). ABP1 showed a positive role for primary root elongation under R at 22 °C, as the *abp1* mutants had shorter roots under the above conditions. Auxin at lower concentrations is known to promote root elongation and inhibits the same at higher concentrations (Ivanchenko et al., 2010). We could complement the root elongation phenotype in Col-0 with the external supplement of auxin in the medium, which was absent in the *abp1* mutant lines indicating that ABP1 binding to auxin might be indispensable for root elongation under R light at 22 °C. Root-localised phytochromes have implications for root elongation (Costigan et al., 2011). We also demonstrated the regulation of PhyB in ABP1-auxin-mediated root development under R at 22 °C. This statement was explained by the observations of drastically diminished or elevated transcript levels of *PHYB* in Col-0 and *abp1* lines respectively. The decrease in root length under R light in *abp1* mutant lines could be due to the elevated *PHYB* levels as compared to Col-0. This observation explained that root elongation is negatively affected due to higher *PHYB* transcript levels in *abp1* mutants. Previous findings of *phyB* levels inhibiting the growth of light-grown roots (Spaninks and Offringa, 2023) supported our results. Our data indicated that ABP1 is involved in decreasing PhyB levels in roots which exerted a positive effect in root elongation under R light at 22 °C. Tissue-specific and pleiotropic developmental role of ABP1 came into the picture when we observed hypersensitivity in hypocotyl length in the case of *abp1* mutants under R light at a lower temperature (18 °C). Here again, regulation by photoreceptor was reclaimed due to higher transcript levels of *PHYB* in the *abp1* mutants. Mutation in *ABP1* resulted in increased *PHYB* levels to exert a positive effect on hypocotyl growth under R light at 18 °C. Hence, it was concluded that ABP1 was involved in decreasing PHYB levels in both roots and hypocotyls to control root elongation and hypocotyl inhibition under R light at 22 °C and 18 °C respectively. The circadian clock coordinates diurnal carbon allocation and controls root growth following a robust diel oscillation of a minimum after (8 h-9 h) dawn and a maximum at the end of the night (Yazdabakhsh et al. 2011). GIGANTEA is not only the component of the night loop of the circadian clock but is also involved in PhyB light signaling for the regulation of flowering time control (Mishra and Panigrahi, 2015). Altered transcript levels of *GI* in the *abp1* mutants indicated a circadian clock control in the root growth phenotype under R at 22 °C. Depleted GI levels in *abp1* mutants indicated its positive role in ABP1-auxin-mediated root growth under R at 22 °C. Thus, ABP1 acts to elevate GI transcript levels to positively affect root elongation under R at 22 °C. Further, ABP1 affected positively the GI transcript levels to control negative hypocotyl growth inhibition under R at 18 °C. Hence, it can be inferred that ABP1 positively affects GI levels to control root and hypocotyl elongation under R light at 22 °C and 18 °C respectively by a yet unknown mechanism.

Interestingly, a positive role of ABP1 in controlling secondary root growth was also observed at 22 °C specifically in B light, where FR light was ineffective and R light was found to have a probable antagonistic role for this B light response. Blue is known to exert additive roles for hypocotyl growth inhibition and root elongation under R light at room temperature (Shinkle et al., 1992; Yeh et al., 2020). Auxin is known to promote lateral root initiation (Ivanchenko et al., 2010). However, the ABP-mediated stimulatory effect on secondary root growth under B light opens up a new area of investigation that requires further experiments including identifying active levels of tissue-specific cryptochromes, phototropins and PhyA. The additive effect of R light to recover the B light phenotype for secondary root growth could be imagined from previous studies (Jeong et al., 2014).

ABP1 function at low temperature was strengthened by observation of delayed flowering phenotype in the *abp1* mutants irrespective of the photoperiod. Corresponding to it decreased levels of GI and FT transcript further supported the delayed flowering in *abp1* mutants. How the ABP1 that is predominantly localized in the ER membrane (>95% at 22 °C) (Oliver et al., 2004; Panigrahi et al., 2009) influences clock output gene (i.e., *GI*) expression needs further investigation. Interestingly, there is no data that shows the intracellular partitioning of ABP1 at low temperatures. This experimental finding along with specific protein interactions in the nucleus if any, would allow to bridge the gap in our understanding between the ABP1 and its role in GI-FT-mediated flowering time control. Notwithstanding, there might be alteration in homeostasis, signaling or sensitivity of other hormones such as cytokinin, auxin or gibberlic acid that may lead to indirect effects on flowering time control at low temperature.

Differential auxin accumulation in the adaxial cells results in leaf epinasty. However, ABP1 binding with auxin has not been conclusive for leaf epinasty (Sandalio et al., 2016). Leaves of *abp1* mutants grown at low temperature showed a more pronounced leaf epinasty phenotype than that of Col-0 indicating that ABP1 has a negative role in leaf epinasty and again reclaiming for its auxin binding at 18 °C. Leaf epinasty can be attributed by various other physiological conditions such as in mutants of phototropins (*phot1phot2*) (Sullivan et al., 2016) and altered ethylene signaling mutants (Barry et al., 2001). Ethylene is known to be synergistic for epinasty in its action with auxin (Mao et al., 2016) to many extents and therefore seems plausible. Investigating the master genes such as Acetyl-CoA synthetase (ACS), 1-Aminocyclopropane-1-Carboxylic Acid Oxidase (ACO), Ethylene Responsive Gene1 (ETR1), Constitutive triple response 1 (CTR1) or downstream regulators such as EINs could also be checked to understand the ABP1-mediated leaf epinasty. Ethylene also is known to orchestrate its action with defence hormones such as SA and JA through ACS, as higher-order *acs* mutants both affect in disease tolerance and also flowering time. Hexaple and heptaple mutants of ACS exhibit earlier flowering phenotype and hypersensitivity to biotrophic pathogens (Putterill et al., 1995). These facts led us to investigate the defence hormone content in the *abp1* mutants. Defence hormones including ABA and SA orchestrated the newer phenotype of lower-temperature tolerance in the *abp1* mutants. An increase of ABA and SA levels in the *abp1* mutants under low temperature when compared to the corresponding Col-0 samples indicated that ABP1 mutation affects levels of these two defence hormones whether via auxin or any unknown pathway. On the other hand, JA levels decreased and followed a contrasting trend to those of ABA and SA. The interplay of these three defence hormones with auxin is plausible along with ethylene and its interaction with ABP1 requires further experiments. Strikingly, the hormonal data suggested that elevated ABA, SA and JA levels were associated with lower-temperature tolerance in the *abp1* mutants.

Cold stress tolerance in the *abp1* mutants was associated with our observations of a decrease in trichome density and an increase in secondary trichome branching (i.e., four-branched trichomes) compared Col-0 under lower-temperature stress. These data suggested that trichome density, which is tightly regulated by cell type-specific abundance *MYB* class transcription factors, was negatively associated and four-branched trichomes were associated positively with the cold stress tolerance in the *abp1* lines. Under lower temperature, the reduced JA levels in the *abp1* mutants could explain the reduced trichome density, as JA significantly is known to promote trichome development by reducing the expression of Jasmonate ZIM-domain (JAZ) proteins eliminating the interactions between JAZ and both bHLH and MYB factors (Fambrini et al., 2019) Under control growth conditions, reduced *CPC* and *TRY* transcript levels explained the dense trichomes in the *abp1* mutants than the Col-0. Moreover, reduced *CPC* transcript levels in Col-0 could correlate to the higher number of trichomes under lower temperature. However, the decreased trichome density in the *abp1* mutants under low temperature could be mediated either due to elevated TRY transcripts in *abp1-C1* or by the other members involved in trichome development (Fambrini et al., 2019), and needs further study. The previous report of positive regulation of auxin for trichome initiation (Wang et al., 2013) is dissimilar to our observation of an increase of trichome density in the *abp1* mutant lines at 22 °C, and requires further investigation. The upregulation of the negative regulator of branching such as *TRY1* is also evidenced by the increased percentage of four-branched trichomes, which may mediate cold stress tolerance, as trichome differentiation was shown to be correlated with cold stress tolerance through TRANSPARENT TESTA GLABRA1 (TTG1), TRIHELIX-TRANSCRIPTION FACTOR GT-2-like 1 (GTL1) genes (Dhawan et al., 2016). Noteworthy that photoreceptors may contribute to cold sensing and trichome branching through their downstream interacting partners i.e., PIF3, PIF4 (Jiang et al., 2017; Pan et al., 2021). Collectively, from the above findings it can be said that abp1 mutation might have some correlation with the trichome density and architecture. Cold stress tolerance may also be an implication of slow dehydration as a whole which has been demonstrated in this study when *abp1* lines showed better tolerance characteristics after 5d of water withheld when compared to those of Col-0 under both 22 °C as well as lower-temperature conditions. The *abp1* mutants showed relatively less water loss, less accumulation of stress metabolite MDA and less enzyme activity of SOD, CAT and POX than the Col-0 lines after WW under both 22 °C as well as low-temperature conditions. The higher dehydration tolerance in the *abp1* mutants could be explained by less ROS (O2^.-^) accumulation probably due to higher carotenoid accumulation, which acts as an efficient quencher for singlet oxygen (Ramel et al., 2012). The non-significant increase in the ROS scavenging enzyme activities in the *abp1* lines with effect to WW may be possible due to the maintenance of higher temperature in these lines as compared to the Col-0 (Figure S4). Hence, it could represent better resistance to the lower temperature of the *abp1* lines than Col-0.

## CONCLUSION

The present study demonstrated the pleiotropic roles of ABP1 in different stages of plant development in selective and specific environmental conditions. The major roles among them are as follows: ABP1 positively regulates root growth under red light, negatively regulates hypocotyl growth inhibition under red light and lower temperature (18 °C) regime and is involved in the regulation of flowering at lower temperature irrespective of photoperiod, in a clock-associated manner and strictly dependent on its binding with auxin. ABP1 bears an antagonistic role in deciphering dehydration tolerance induced due to lower temperature or water withdrawal of 5 days. ABP1 mutation is associated with altered the phytohormone levels including abscisic acid, jasmonic acid and salicylic acid. Dehydration tolerance in *abp1-C1* and *abp1-C2* mutant lines is suggestively attained by altered defense hormone levels, trichome density and architecture. These new findings have opened novel avenues for functional validations to understand the molecular mechanisms of these processes mediated by the ABP1.

## ACKNOWLEDGEMENT

We are grateful to the Department of Atomic Energy, Government of India and NISER for providing the experimental facility at the institute. We thank Prof. Dr. Mark Estelle, University of California, San Diago, Dr. Jiri Friml, Institute of Science and Technology, Austria and Dr. Stefan Kircher, University of Freiburg, Germany for their generous support in providing *abp1-C1* and *abp1-C2* seeds respectively for our research. Furthermore, we would also like to thank the Metabolomics facility at NIPGR for carrying out the defence hormone quantification.

## AUTHORS CONTRIBUTIONS

The experiments were conceived and designed by KCP and AP. The experiments were carried out by AP, AB, AK and AD. The preliminary work was established by NP and MS. AP, MP and KCP wrote and revised the manuscript and analyzed the data. The manuscript has been read and approved by all the authors.

---

**Figure S1.**
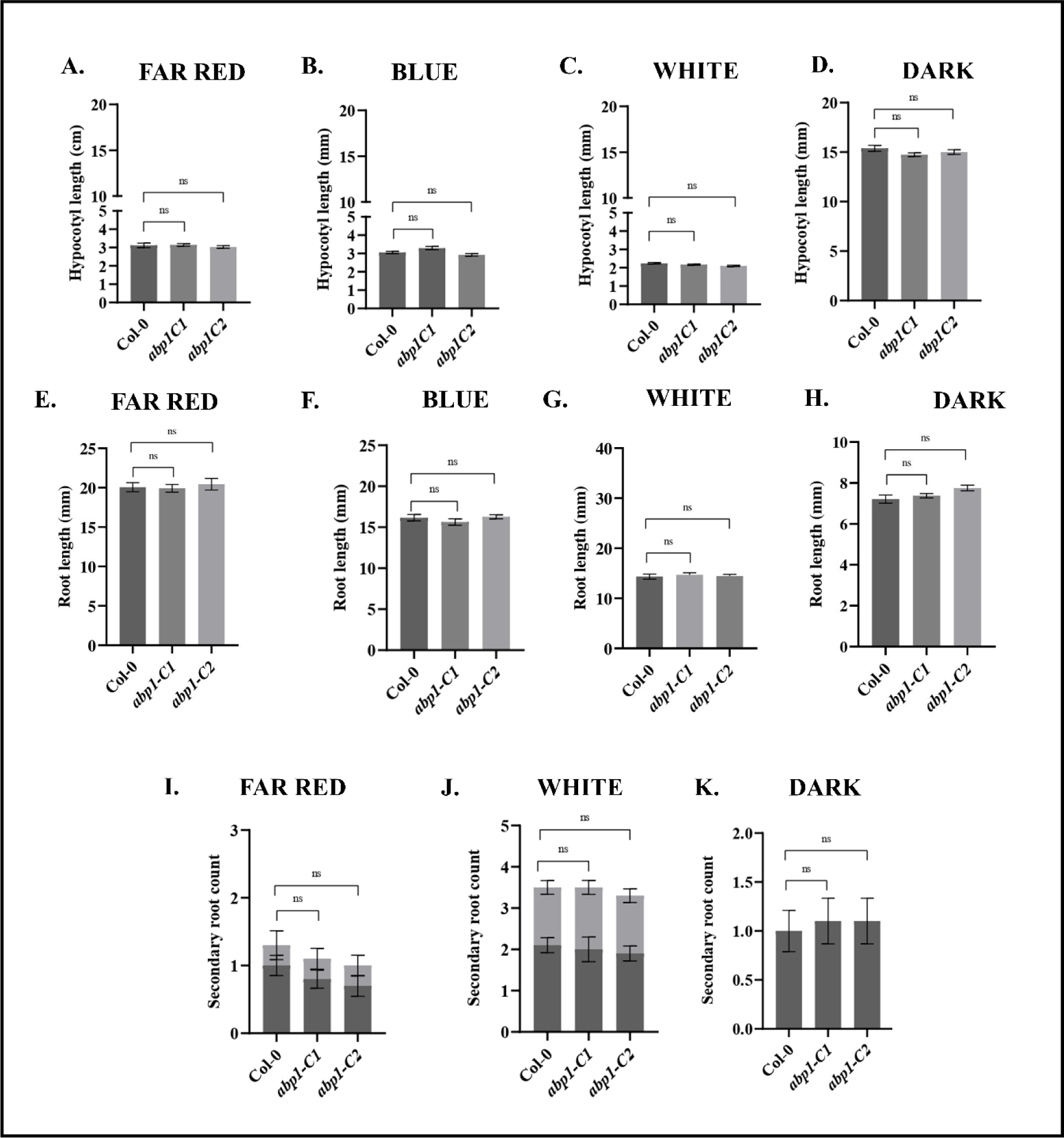
Seedling phenotypes were non-significant under different monochromatic lights at 22 °C. Seedlings of Col-0, *abp1-C1* and *abp1-C2* lines were grown on MS media under LD conditions and different monochromatic lights of Far-red (**A, E, I**), Blue (**B, F**), White (**C, G, J**) or in darkness (**D, H, K**) till 7-days, after which hypocotyl growth (**A-D**) or root length (**E-H**) and secondary root counts (**I-K**) were measured. Nearly 300 seedlings were grown in 3 biological replicates with 100 seedlings each time. Data presented as mean ± SE. One-way analysis of variance (ANOVA) using Tukey’s multiple comparisons test was performed with the help of GraphPad Prism to test for significance among the dataset, \**p* < 0.05, **< 0.01, ns – non-significant.

**Figure S2.**
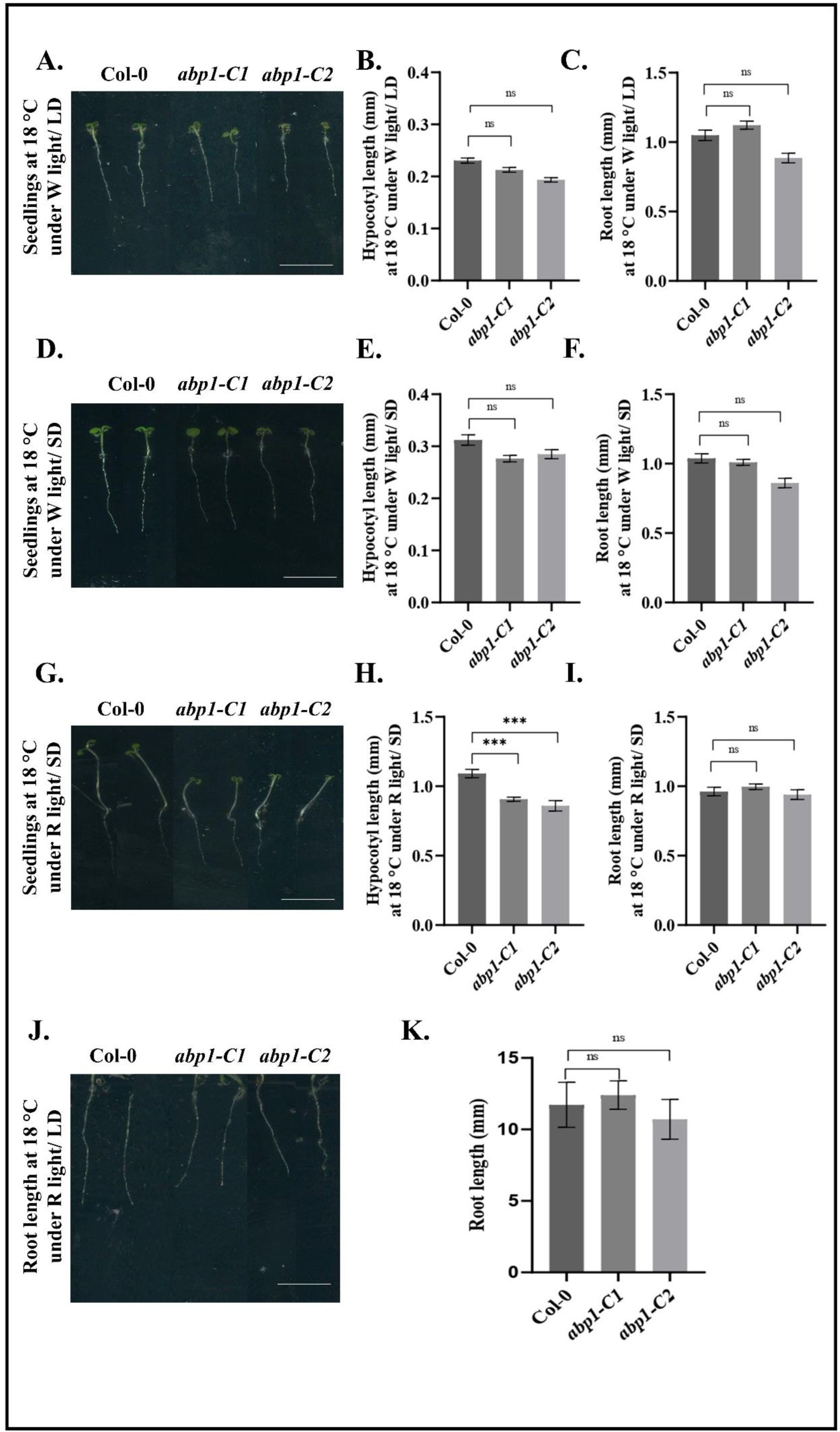
Seedling phenotype at lower temperature i.e., 18 °C. Seedlings of Col-0, *abp1-C1* and *abp1-C2* lines were grown on MS media under LD (**A-C**), SD (**D-F**) conditions under W light and SD (**G-I**), LD (**J-K**) under R light at 18 °C. Hypocotyl and root length were measured using ImageJ software. Nearly 300 seedlings were grown in 3 biological replicates with 100 seedlings each time. Data presented as mean ± SE. One-way analysis of variance (ANOVA) using Tukey’s multiple comparisons test was performed with the help of GraphPad Prism to test for significance among the dataset, \**p* < 0.05, **< 0.01, ns – non significant. Scale bar indicates 5 mm.

**Figure S3.**
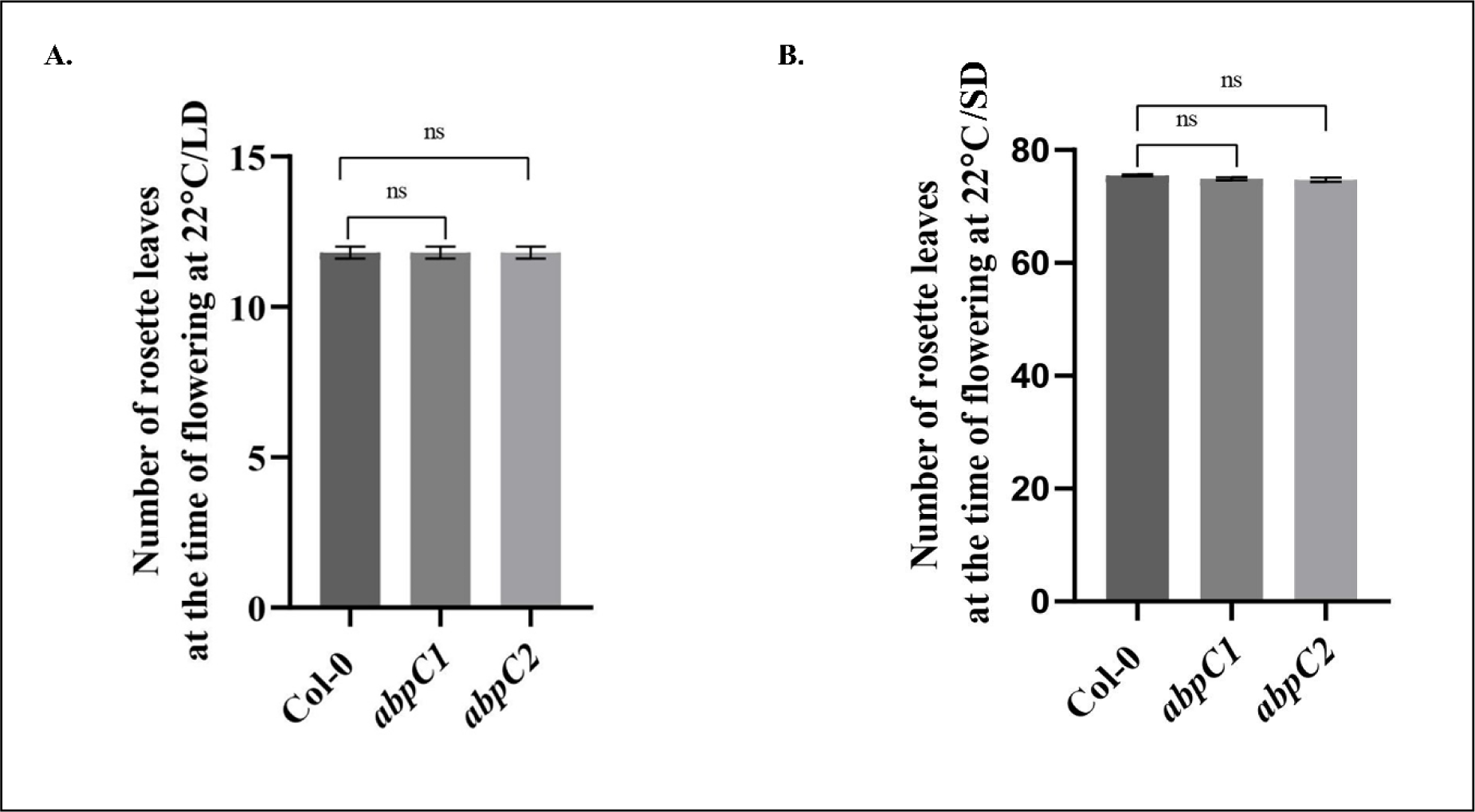
Flowering phenotype under LD and SD conditions at 22 °C. Plants of Col-0, *abp1-C1* and *abp1-*C2 genotypes were grown on soil in pots under LD or SD conditions at 22 °C till bolting. The total number of rosettes leaves at the time of bolting grown under LD (**A**) and SD (**B**) conditions under 22 °C were noted. Three biological replicates were performed using 10 plants in each replicate. Data presented as mean ± SE. One-way analysis of variance (ANOVA) using Tukey’s multiple comparisons test was performed with the help of GraphPad Prism to test for significance among the dataset, * *p* < 0.05, ** < 0.01, *** *p* < 0.001and ns represent non-significant differences.

**Figure S4.**
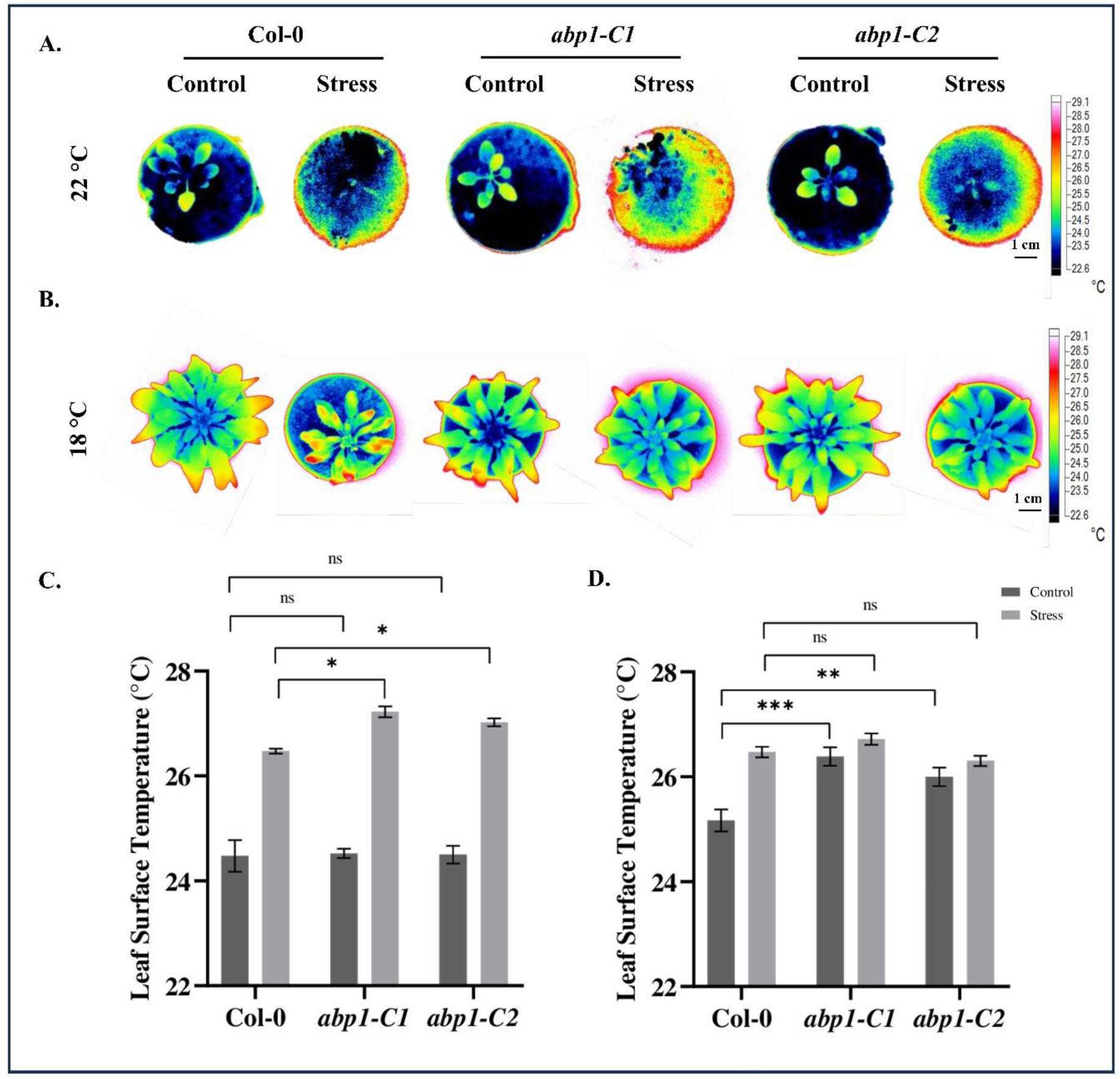
Plant leaf Temperature analysis. **A and B.** Plants of Col-0, *abp1-C1* and *abp1-* C2 genotypes were grown on soil in pots under LD conditions at 22 °C (**A**) or 18 °C (**B**) till bolting stage. Plants were either well-watered or water withheld for 5-10 days (WW) as described in materials and methods. Leaf temperature of the plants grown in water withheld conditions at 22 °C (**A and C**) or 18 °C (**B and D**) were measured from thermal pictures that were taken with a FLUKE infrared camera and analyzed using the SMART VIEW software. Leaf surface temperature showing highest temperatures were selectively chosen from at least ten individual plants. Three individual biological replicates were performed. Data presented as mean ± SE. One-way analysis of variance (ANOVA) using Tukey’s multiple comparisons test was performed with the help of GraphPad Prism to test for significance among the dataset, * *p* < 0.05, ** < 0.01, *** *p* < 0.001and ns represent non-significant differences. Scale bar: 1 cm.

**Table S1.**
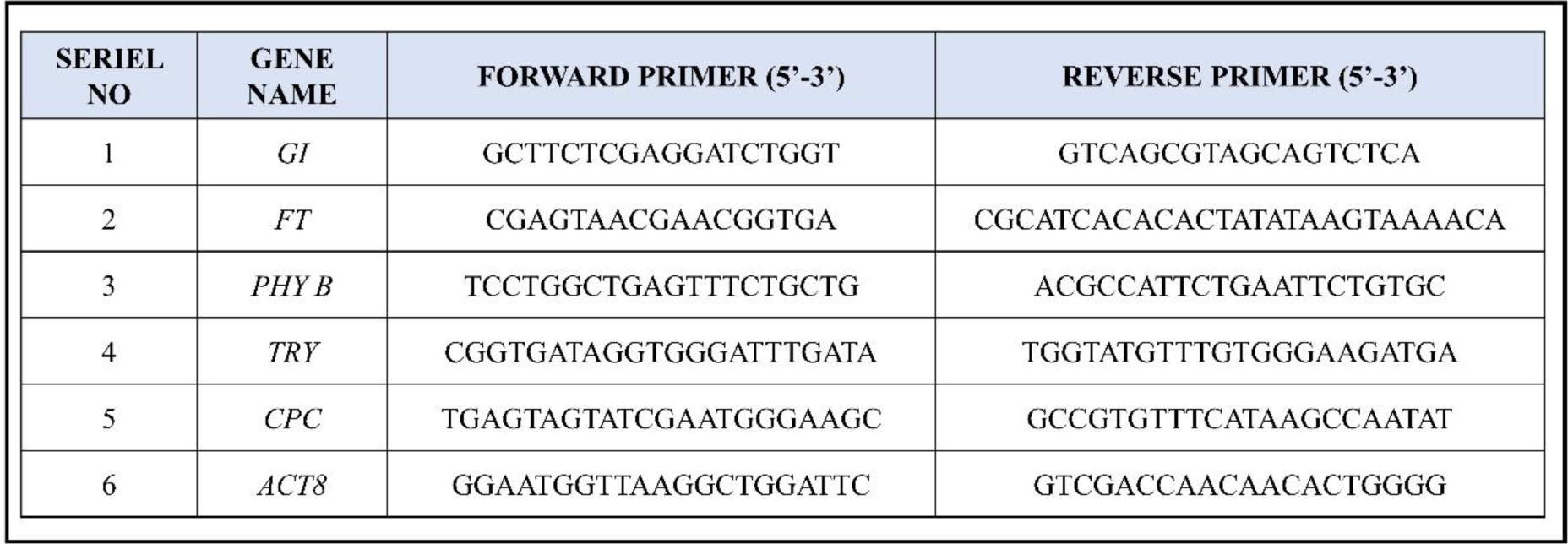
List of primers used for q-PCR.

